# Clinical Utility of Expanded Carrier Screening: Reproductive Behaviors of At-Risk Couples

**DOI:** 10.1101/069393

**Authors:** Caroline Ghiossi, James D. Goldberg, Imran S. Haque, Gabriel A. Lazarin, Kenny K. Wong

## Abstract

**Purpose:** Expanded carrier screening (ECS) analyzes dozens or hundreds of recessive genes for determining reproductive risk. Data on clinical utility of screening conditions beyond professional guidelines is scarce.

**Methods:** Individuals underwent ECS for up to 110 genes. 537 at-risk couples (ARC), those in which both partners carry the same recessive disease, were invited to a retrospective IRB-approved survey of their reproductive decision making after receiving ECS results.

**Results:** 64 eligible ARC completed the survey. Of 45 respondents screened preconceptionally, 62% (n=28) planned IVF with PGD or prenatal diagnosis (PNDx) in future pregnancies. 29% (n=13) were not planning to alter reproductive decisions. The remaining 9% (n=4) of responses were unclear.

Of 19 pregnant respondents, 42% (n=8) elected PNDx, 11% (n=2) planned amniocentesis but miscarried, and 47% (n=9) considered the condition insufficiently severe to warrant invasive testing. Of the 8 pregnancies that underwent PNDx, 5 were unaffected and 3 were affected. 2 of 3 affected pregnancies were terminated.

Disease severity was found to have significant association (p=0.000145) with changes in decision making, whereas guideline status of diseases, controlled for severity, was not (p=0.284).

**Conclusion:** Most ARC altered reproductive planning, demonstrating the clinical utility of ECS. Severity of conditions factored into decision making.

## INTRODUCTION

Carrier screening identifies couples at increased risk of having a child with a genetic disease and enables them to consider alternative reproductive options. Those who do not make alternative reproductive decisions based on their carrier status may still use this knowledge to prepare for the birth of an affected child and facilitate early intervention (in many cases) for the best possible outcomes.^1,2^

Historically, carrier screening programs targeted a small number of diseases that are highly prevalent in an ethnic-defined population. More recently developed, expanded carrier screening (ECS) assesses risk for dozens or hundreds of diseases across all populations (panethnic, or universal screening).^3^ Despite widespread adoption of ECS by many providers, statements from professional organizations suggest additional research is needed.^3,4^ In particular, they cite the lack of data regarding reproductive outcomes of couples that undergo ECS. Studies of outcomes after population-based carrier screening initiatives for a limited number of disorders have consistently found a reduced incidence of the disease of interest due to the decisions made by the at-risk couples (ARC). Tay-Sachs disease incidence fell by 90% in the Ashkenazi Jewish population due to high screening uptake over the course of multiples decades.^5^ Screening for thalassemia in Mediterranean and Chinese populations has resulted in similar declines.^5,6,7^

Decision making among couples at risk for children affected by cystic fibrosis (CF) has been assessed in several studies. Many examined pregnant couples, often focusing on those who have an affected child or relative. While these studies indicated that the majority of couples would pursue prenatal diagnosis in a pregnancy ^8,9,10^, having an affected child or relative may influence reproductive decision making; while decisions of carrier couples in this context are well characterized, they do not represent the majority experience of at-risk carrier couples.^9^ The available studies that are focused on CF screening in the general population, not based on family history or known carrier status, do demonstrate effectiveness in reducing the incidence of the condition. In the United States, a study in Massachusetts demonstrated a decrease in CF incidence following the preconception and prenatal screening recommendations by ACOG, ACMG and the National Institutes of Health.^11^ Several studies have been conducted outside of the US; Of significance is a 5-year study in Edinburgh, UK that showed a drop in CF incidence after the implementation of a CF prenatal screening program.^12^ A 3-year study of a screening program in Australia identified 9 ARC, 6 that were pregnant at the time of screening. All 6 couples elected prenatal diagnosis, and 3 preconception couples elected *in vitro* fertilization (IVF) with preimplantation genetic diagnosis (PGD) in future pregnancies.^13^

Literature regarding the clinical utility of ECS panels is only beginning to emerge. One recent study by Franasiak et al. focused on the clinical decision making of infertile couples found to be at-risk carriers through ECS testing as part of their fertility work-up. In total 8 couples were identified as at-risk carriers and all elected to pursue PGD as part of their IVF treatment, indicating that the results of the carrier screening affected clinical decision making in all cases, though the authors note that not all of the couples followed through with their planned treatment.^14^

The purpose of this study is to learn about the reproductive decisions of ARC as identified by ECS from a nationwide population. This paper will describe the experience of ARC after they received their ECS results, characterize their reproductive decisions, and identify factors associated with their decision making.

## MATERIALS AND METHODS

### Eligibility Criteria

This was a mixed-methods retrospective study in which participants were invited to self-report their experience and outcomes. The study was approved by the California State University Stanislaus Institutional Review Board, Protocol #1516-007.

Participants were selected from those receiving Expanded Carrier Screening (Family Prep Screen 1.0 or Family Prep Screen 2.0) through Counsyl (South San Francisco, CA), a molecular diagnostics laboratory. The Family Prep Screen tests for carrier status in up to 110 genes, either by targeting 417 predefined disease-causing mutations (version 1.0) or by next-generation exonic sequencing and pathogenicity interpretation for novel sequence variants (version 2.0). Test orders required physician authorization, typically a specialist in obstetrics, reproductive endocrinology, maternal-fetal medicine, or clinical genetics. Carrier screening was voluntary and consent to research was included within the general consent form, with the option to request exclusion. Genetic counseling was made available at no additional cost to all individuals tested.

Conditions included in the ECS panel range in severity. A method for severity classification divided diseases into four groups, from most to least impactful: profound, severe, moderate, and mild ^15^. In this classification algorithm, disease characteristics are organized into tiers based on their impact to the affected individual. Severity is assigned to a disease based a combination of the number of characteristics present in disease, and the tier ranking of those characteristics ^15^.

At-risk couples (ARC) were defined as self-identified reproductive partners in which both were identified as carriers for the same profound, severe, or moderate autosomal recessive condition. ARC identified by the laboratory between April 2014 and August 2015 were selected for inclusion in the study if contact information was available, neither individual had requested exclusion from research, and neither known personal carrier status of genetic disease nor testing for gamete donor candidacy was reported on the test requisition.

Couples at risk for a mild condition (e.g. pseudocholinesterase deficiency, OMIM +177400) or couples in which the female carried an X-linked condition (e.g. fragile X, OMIM #300624) were excluded. The former was decided in order to focus the study on diseases with greatest clinical impact, while the latter was meant to ensure that the risk for an affected child was consistent across couples, for autosomal recessive diseases, 25%.

### Survey Design

The survey comprised 33 questions and was deployed online through SurveyGizmo (Boulder, CO), a survey research tool providing HIPAA-compliant privacy and security features. The survey included branch logic to skip or display certain questions based on each individual’s answer about their pregnancy status and reproductive decisions. The survey questions were designed to elicit ARC’s experience with ECS, their reproductive choices, and future plans after receiving their ECS results. Choices assessed included *in vitro* fertilization (IVF) with preimplantation genetic diagnosis (PGD), prenatal diagnosis such as chorionic villus sampling (CVS) or amniocentesis, termination of pregnancy, adoption, gamete donation, no longer planning to have children, or (as free text) other changes in reproductive plans. Additionally, respondents could indicate that they were not planning to pursue any alternative options. Questions also included the condition for which they were found to be carriers, reasons for pursuing carrier screening, utilization of and satisfaction with genetic counseling, pregnancy history, and demographics. The survey consisted of multiple choice questions to capture categorical variables as quantitative data, as well as open-ended questions about participants’ experiences. Responses to the survey were anonymous.

The full survey is provided in the supplemental information.

### Data Collection

Data were collected from two sets of ARC. The first set (n=465) were first invited by email, followed by a reminder email and/or a SMS text. A second set (n=72) without email or mobile phone contact information on file were invited via paper mail. Upon survey completion, participants were given the option to enter their email address in a drawing for Amazon.com gift cards.

### Data Analysis

Survey data was analyzed in SPSS (IBM Corporation, Armonk, NY). Descriptive statistics were generated to characterize the trends in the data. Data categories were collapsed to 2x2 contingency tables and analyzed using Fisher’s exact test. To control for multiple hypothesis testing, the significant p-value was adjusted using the Bonferroni correction.

Free text responses were analyzed in Excel (Microsoft Corporation, Redmond, WA). An open coding framework was used to identify broad themes in the participants’ experiences, decision-making rationale, and future plans. This qualitative data was used to enrich the quantitative analysis and provide patient perspective and commentary on the trends seen in the statistics.

## RESULTS

In total, the eligibility criteria yielded 537 ARC for possible participation. Of those, 465 could be contacted by SMS or email, and the remaining 72 could only be contacted through postal mail.

The online survey received a 18% (n=86) total response rate including 16 partial responses in which the respondent either declined consent and was exited from the survey, or provided consent but exited the survey prior to reaching the end. 15% (n=70) completed the survey. Of the 72 paper surveys mailed via post, 13% (n=9) were returned completed. Completion rate was not significantly different between methods (p=0.72).

Of 79 completed surveys, 3 were excluded because the respondents did not report the condition for which they were both carriers, instead choosing either “I don’t recall” or “My partner and I were not carriers for the same condition.” An additional 12 that reported a family history of the condition for which they were found to be carriers were excluded from this analysis.

The demographic data of the remaining 64 participants are summarized in Table 1. Respondents and their partners were predominantly Caucasian (76% and 70% respectively selecting this option), educated (89% and 77% respectively with bachelor′s degree or higher), with annual household incomes exceeding $100,000 (68%). The majority of female partners were 25-34 years of age (81%), had no children at the time of responding to the survey (62%), had no history of miscarriage (69%). Most ARC had carrier screening as part of a fertility work-up (53%) with other reasons for carrier screening including routine screening (31%), ethnicity-based screening (6.3%), prior miscarriages or stillbirth (4.7%), ultrasound anomalies (3.1%), and consanguinity (1.6%). ARC reported having ECS on the recommendation of a healthcare provider (86%) and almost all pursued genetic counseling after receiving results (95%), the majority of those through Counsyl’s services (61%).

**Figure 1.**
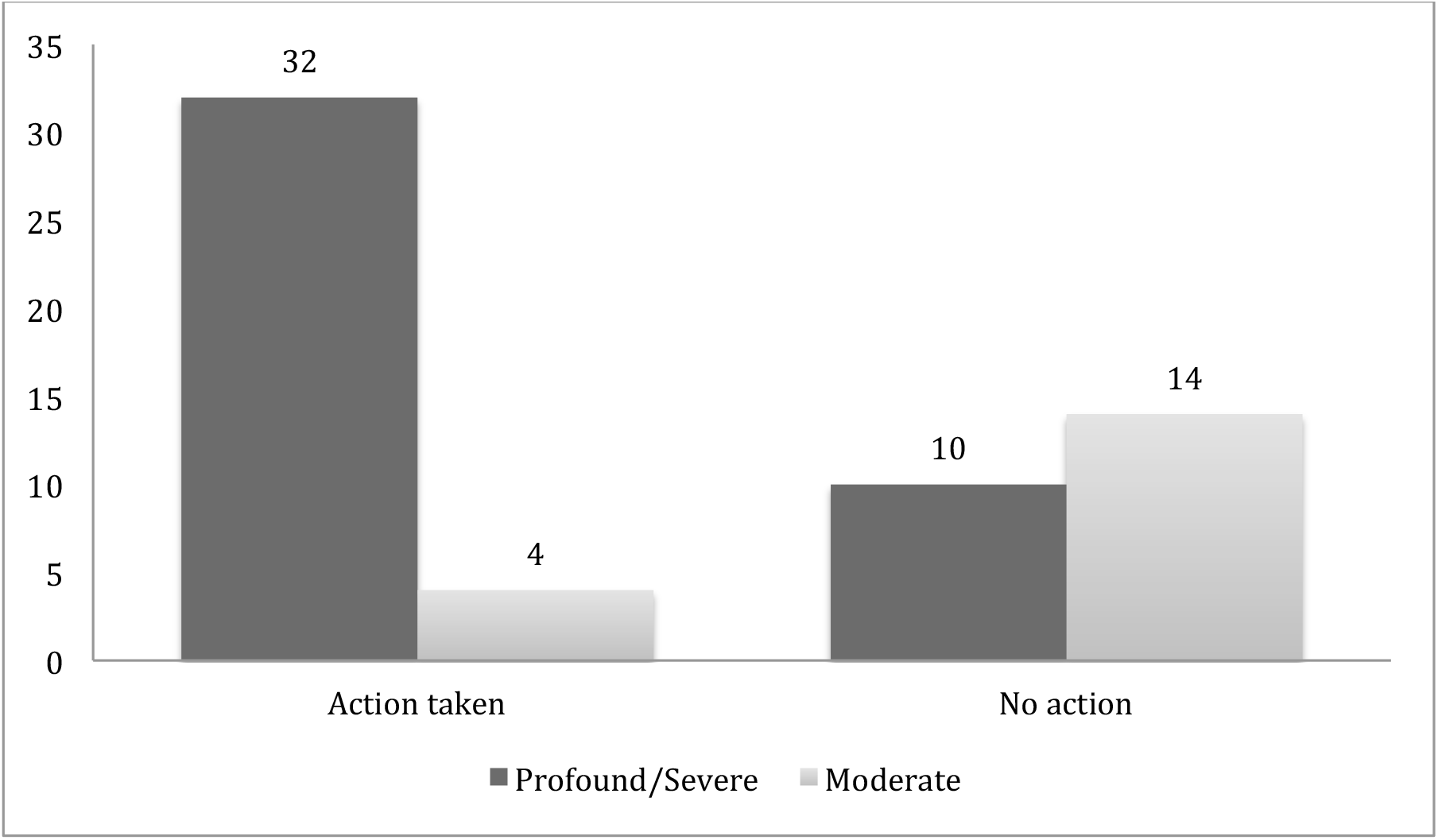
Disease severity and action taken/planned based on ECS results. Note: Biotinidase deficiency (BTD) is classified as a severe condition in this analysis. However, there is a wide spectrum of severity in BTD symptoms with partial deficiencies that lead to a mild - moderate presentation.

**Table 1.** Participant demographic information

| Characteristic | Total Respondents | Categories | Responses | Percentage |
| --- | --- | --- | --- | --- |
| **Female partner's age** | 64 | 18 – 24 | 2 | 3.1% |
|  |  | 25 – 34 | 52 | 81% |
|  |  | 35 – 44 | 10 | 16% |
| **Number of children** | 63 | None | 39 | 62% |
|  |  | 1 | 21 | 33% |
|  |  | 2+ | 3 | 4.8% |
| **Past miscarriages** | 64 | Yes | 20 | 31% |
|  |  | None | 44 | 69% |
| **Ethnicity** | 59 (*multiple selections allowed*) | European/Mixed Caucasian | 45 | 66% |
|  |  | Ashkenazi Jewish | 11 | 16% |
|  |  | Asian | 4 | 5.9% |
|  |  | African American | 3 | 4.4% |
|  |  | Others | 5 | 7.4% |
| **Partner's ethnicity** | 58 (*multiple selections allowed*) | European/Mixed Caucasian | 41 | 64% |
|  |  | Ashkenazi Jewish | 12 | 19% |
|  |  | Asian | 4 | 6.3% |
|  |  | African American | 3 | 4.7% |
|  |  | Others | 4 | 6.3% |
| **Religious affiliation** | 64 | Protestant | 13 | 20% |
|  |  | Catholic | 14 | 22% |
|  |  | Jewish | 11 | 17% |
|  |  | Others | 5 | 7.8% |
|  |  | None | 17 | 27% |
|  |  | Decline to state | 4 | 6.3% |
| **Partner's religious affiliation** | 63 | Protestant | 12 | 19% |
|  |  | Catholic | 9 | 14% |
|  |  | Jewish | 12 | 19% |
|  |  | Others | 3 | 4.8% |
|  |  | None | 23 | 37% |
|  |  | Decline to state | 4 | 6.3% |
| **Highest level of education** | 64 | High school/ vocational school | 3 | 4.7% |
|  |  | Some college/ associate degree | 4 | 6.3% |
|  |  | Bachelor degree | 23 | 36% |
|  |  | Graduate degree | 34 | 53% |
| **Partner's highest level of education** | 64 | High school/ vocational school | 5 | 7.8% |
|  |  | Some college/ associate degree | 10 | 16% |
|  |  | Bachelor degree | 19 | 30% |
|  |  | Graduate degree | 30 | 47% |
| **Annual household income** | 63 | <\$49,999 | 7 | 11% |
| | | \$50,000 - \$74,999 | 5 | 7.9% |
| | | \$75,000 - \$99,999 | 8 | 13% |
| | | \$100,000 - \$150,000 | 23 | 37% |
| | | >\$150,000 | 20 | 32% |

**Table 2.** Motivations and circumstances for pursuing carrier screening

| **Characteristic** | **Total respondents** | **Categories** | **Responses** | **Percentage** |
| --- | --- | --- | --- | --- |
| **Reason for screening** | 64 | Fertility work-up | 34 | 53% |
|  |  | Routine screening | 20 | 31% |
|  |  | Multiple miscarriages/stillbirth | 3 | 4.7% |
|  |  | Ethnicity | 4 | 6.3% |
|  |  | Consanguinity | 1 | 1.6% |
|  |  | Ultrasound anomaly | 2 | 3.1% |
| **Initiator of screening** | 63 | Healthcare provider | 54 | 86% |
|  |  | Patient requested | 9 | 14% |
| **Length of time since receiving results** | 64 | 1-3 months | 11 | 17% |
|  |  | 3-6 months | 13 | 20% |
|  |  | 6-9 months | 8 | 13% |
|  |  | >9 months | 32 | 50% |
| **Genetic counseling (GC) services** | 64 | Counsyl GC | 37 | 58% |
|  |  | Local GC | 22 | 34% |
|  |  | Other provider | 2 | 3.1% |
|  |  | None | 3 | 4.7% |
| **Disease severity classification** | 64 | Moderate | 19 | 30% |
|  |  | Severe | 34 | 53% |
|  |  | Profound | 11 | 17% |
| **Pregnancy status at time of receiving results** | 64 | Pregnant | 19 | 30% |
|  |  | Preconception | 45 | 70% |

60 participants provided an interpretable answer about their actions or planned actions following results receipt. The remaining 4 participants either declined to answer the question or did not supply an interpretable response (e.g. selecting “other” without additional explanation when asked about their plans or selecting/indicating contradictory options.) The reproductive decisions reported by ARC were collapsed into two categories: action (36/60) or no action (24/60) based on the ECS results, with the unclear responses (4/64) excluded from the statistical analysis. Actions reported by this group included IVF with PGD (n=22) and prenatal diagnosis (n=14).

Potential associations between alternative reproductive options and disease severity, pregnancy status (non-pregnant ARC have more available options), and demographic factors were assessed (Table 3). After Bonferroni correction, the required threshold for significance was .05/10 = *α* =.005.

**Table 3.**
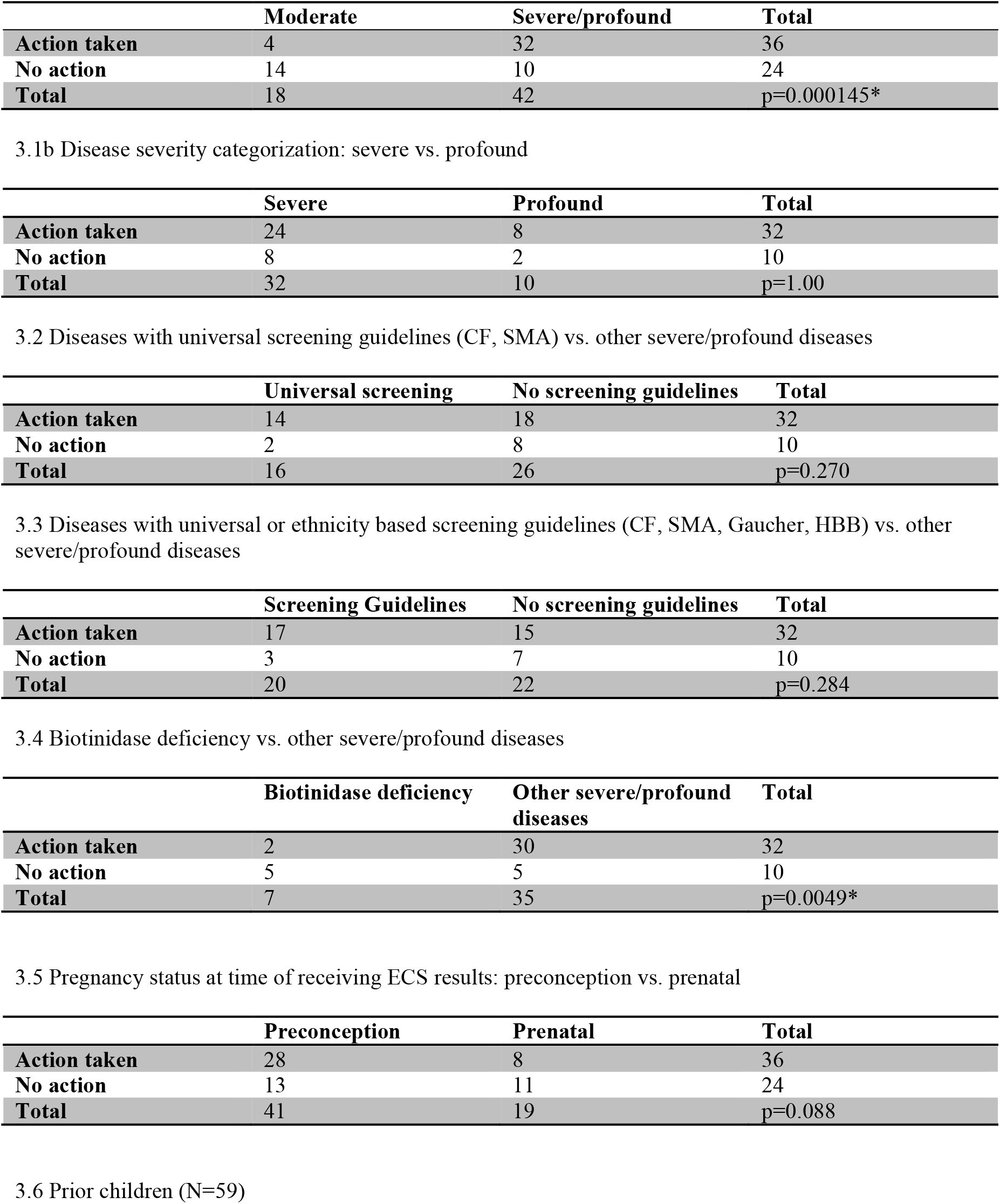
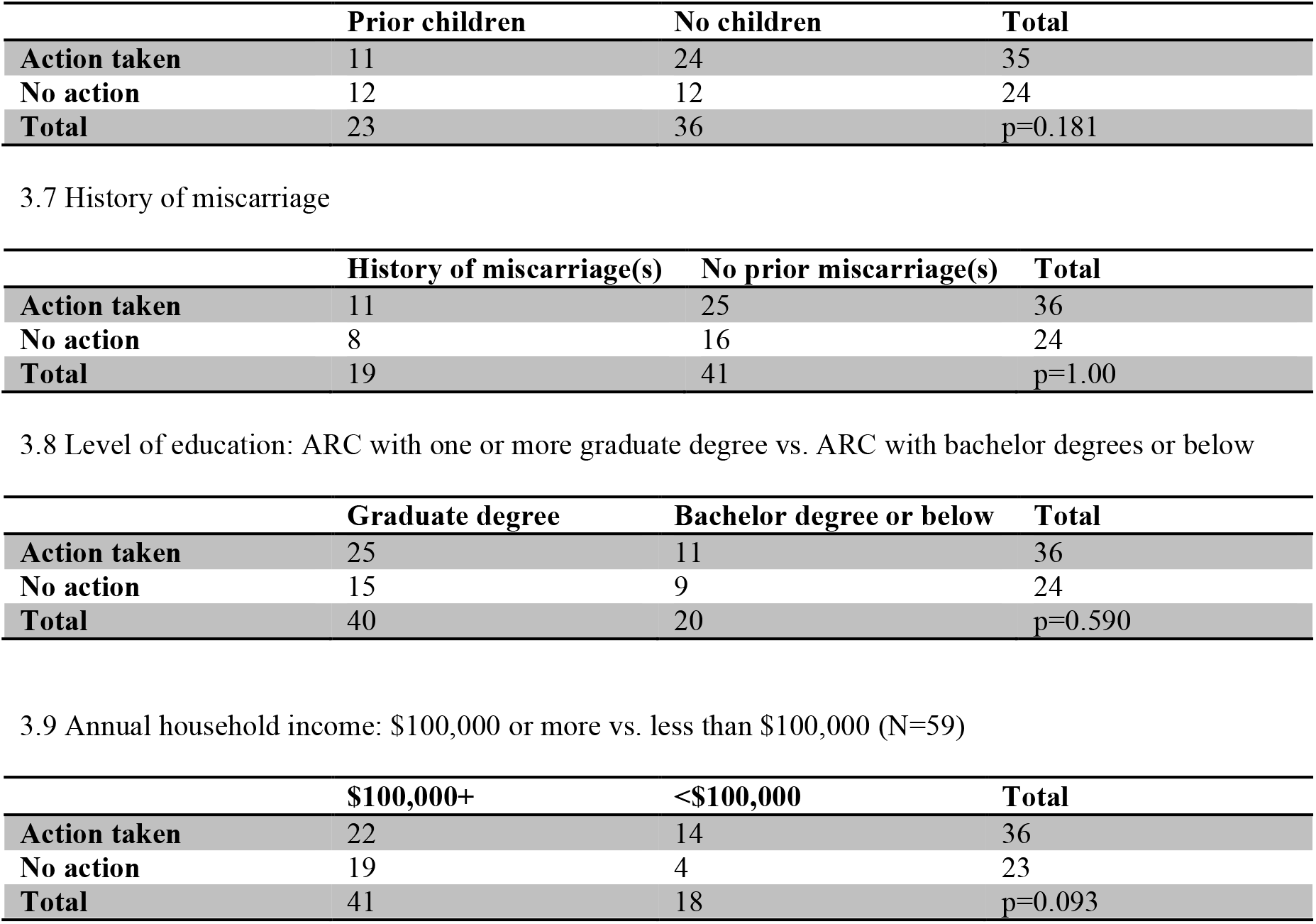
Associations with reproductive decisions. Data on these variables were collapsed into a 2x2 contingency table and analyzed using Fisher’s exact. Demographics and fertility history were collapsed to the majority category versus all other categories as described below. Action taken based on ECS results was collapsed to action taken or no action.

3.1a Disease severity categorization: moderate vs. severe/profound
|  | **Moderate** | **Severe/profound** | **Total** |
| --- | --- | --- | --- |
| **Action taken** | 4 | 32 | 36 |
| **No action** | 14 | 10 | 24 |
| **Total** | 18 | 42 | p=0.000145* |

3.1b Disease severity categorization: severe vs. profound
|  | **Severe** | **Profound** | **Total** |
| --- | --- | --- | --- |
| **Action taken** | 24 | 8 | 32 |
| **No action** | 8 | 2 | 10 |
| **Total** | 32 | 10 | p=1.00 |

3.2 Diseases with universal screening guidelines (CF, SMA) vs. other severe/profound diseases
|  | **Universal screening** | **No screening guidelines** | **Total** |
| --- | --- | --- | --- |
| **Action taken** | 14 | 18 | 32 |
| **No action** | 2 | 8 | 10 |
| **Total** | 16 | 26 | p=0.270 |

3.3 Diseases with universal or ethnicity based screening guidelines (CF, SMA, Gaucher, HBB) vs. other severe/profound diseases
|  | **Screening Guidelines** | **No screening guidelines** | **Total** |
| --- | --- | --- | --- |
| **Action taken** | 17 | 15 | 32 |
| **No action** | 3 | 7 | 10 |
| **Total** | 20 | 22 | p=0.284 |

3.4 Biotinidase deficiency vs. other severe/profound diseases
|  | **Biotinidase deficiency** | **Other severe/profound diseases** | **Total** |
| --- | --- | --- | --- |
| **Action taken** | 2 | 30 | 32 |
| **No action** | 5 | 5 | 10 |
| **Total** | 7 | 35 | p=0.0049* |

3.5 Pregnancy status at time of receiving ECS results: preconception vs. prenatal
|  | **Preconception** | **Prenatal** | **Total** |
| --- | --- | --- | --- |
| **Action taken** | 28 | 8 | 36 |
| **No action** | 13 | 11 | 24 |
| **Total** | 41 | 19 | p=0.088 |

3.6 Prior children (N=59)
|  | **Prior children** | **No children** | **Total** |
| --- | --- | --- | --- |
| **Action taken** | 11 | 24 | 35 |
| **No action** | 12 | 12 | 24 |
| **Total** | 23 | 36 | p=0.181 |

3.7 History of miscarriage
|  | **History of miscarriage(s)** | **No prior miscarriage(s)** | **Total** |
| --- | --- | --- | --- |
| **Action taken** | 11 | 25 | 36 |
| **No action** | 8 | 16 | 24 |
| **Total** | 19 | 41 | p=1.00 |

3.8 Level of education: ARC with one or more graduate degree vs. ARC with bachelor degrees or below
|  | **Graduate degree** | **Bachelor degree or below** | **Total** |
| --- | --- | --- | --- |
| **Action taken** | 25 | 11 | 36 |
| **No action** | 15 | 9 | 24 |
| **Total** | 40 | 20 | p=0.590 |

3.9 Annual household income: \$100,000 or more vs. less than \$100,000 (N=59)
| | **\$100,000+** | **<\$100,000** | **Total** |
| --- | --- | --- | --- |
| **Action taken** | 22 | 14 | 36 |
| **No action** | 19 | 4 | 23 |
| **Total** | 41 | 18 | p=0.093 |

Of ARC carrying severe or profound conditions, 76% (32/42) reported alternative reproductive actions, versus 22% (4/18) ARC carrying moderate conditions, suggesting that disease severity has a significant effect on reproductive actions (p=0.000145). One severe condition was a clear outlier to this trend: only 29% (2/7) of ARC carrying biotinidase deficiency reporting a change in their actions, in contrast to the 86% (30/35) of ARC carrying other severe or profound conditions (p=0.0049). Of ARC carrying profound conditions, 80% (8/10) reported alternative reproductive actions vs. 75% (24/32) of ARC carrying severe conditions (p=1.00).

Because ECS includes all of the conditions recommended by ACOG and ACMG, the data allow us to compare ARC actions between those diseases with guidelines and those without. For diseases recommended for panethnic screening (CF and SMA), 14 ARC reported taking action on the results while 2 did not alter their reproductive plans. There was no significant difference (p=0.270) between rate of action on these recommended conditions as compared to the other severe or profound conditions with 18 in that group reporting action based on the result and 8 reporting no change to reproductive plans (Table 4). If all diseases that are included in guidelines, including those for ethnicity based screening, are compared, 17 reported taking action based on the result, while 3 did not. Here again, there was no significant difference (p=0.284) between ARC carrying diseases with screening recommendations and the ARC carrying non-guideline diseases with 15 reporting action taken based on their results and 7 reporting no change in reproductive plans.

**Table 4.**
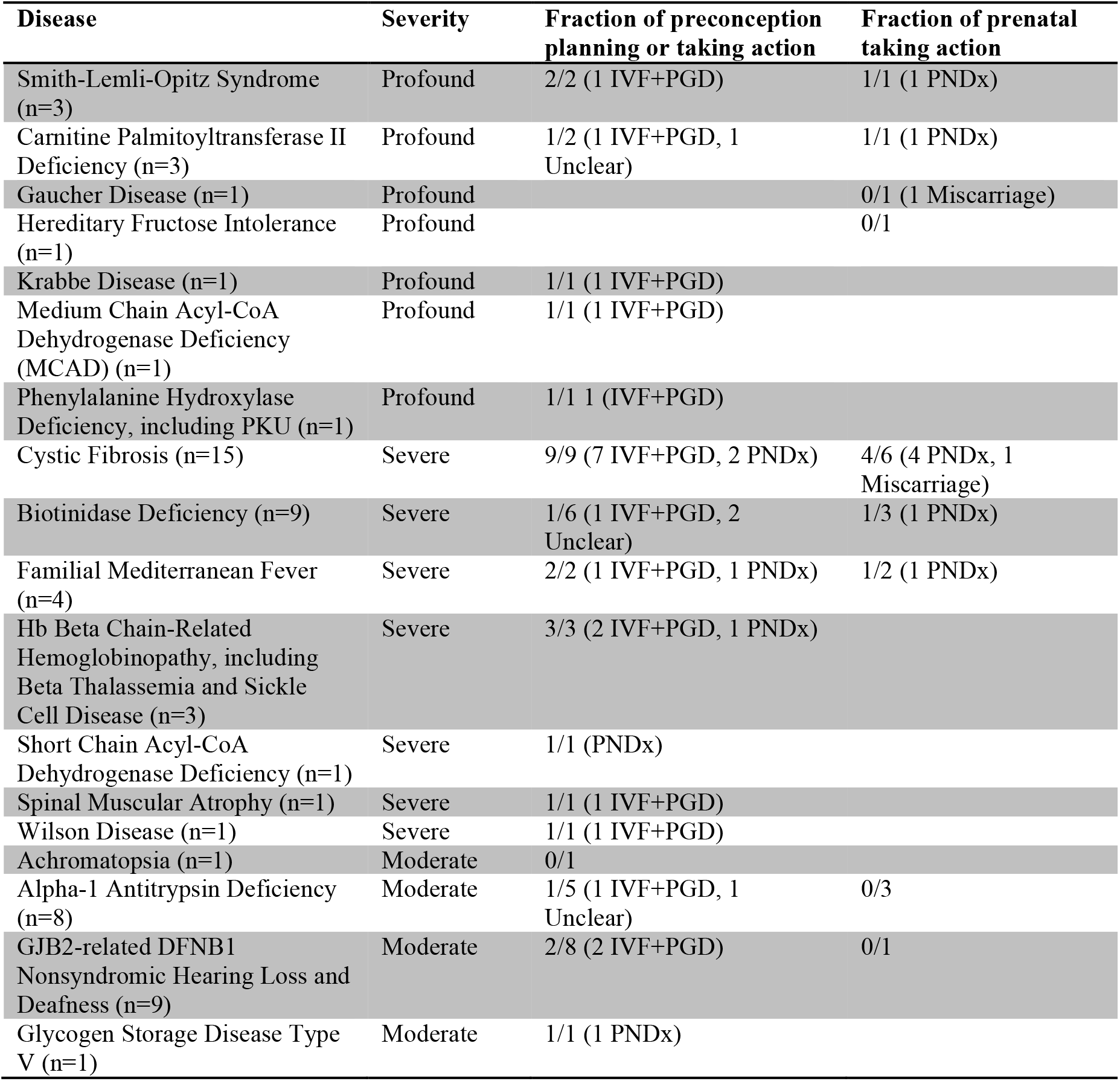
Diseases and corresponding reproductive decisions in the preconception and prenatal contexts(N=64)

In contrast to the hypothesis that the additional options conferred by preconception screening would lead to more alternative actions taken by non-pregnant couples, pregnancy status was not found to be a significant variable (p=0.088). Existence of prior children (p=0.181), occurrence of prior miscarriages (p=1.00), attainment of graduate education by at least one member of the couple (p=0.590), and annual income greater than or equal to US$100,000 (p=0.093) were factors not found to have significant associations.

### Actions in the Prenatal Context

19 of 64 (30%) ARC were pregnant when they received ECS results. Of these, 42% (8/19) elected prenatal diagnosis (CVS or amniocentesis) and 58% (11/19) did not. Of the latter group, 2 participants reported planning a diagnostic procedure but miscarried before the procedure could be done, effectively bringing those who took action or planned to take action to 53% (10/19) and those who did not plan to take action to 47% (9/19). Both of the ARC who experienced a miscarriage prior to planned prenatal diagnosis indicated that they would pursue IVF with PGD in future pregnancies. Of the 8 ARC who underwent prenatal diagnosis, 5 fetuses did not inherit both parental mutations and 3 were homozygous or compound heterozygous for the parental mutations, consistent with Hardy-Weinberg expectations (p=0.422, exact binomial test). Two of the latter three pregnancies were voluntarily terminated (carnitine palmitoyltransferase II deficiency OMIM *600650 and cystic fibrosis OMIM #219700) and one was continued (cystic fibrosis).

### Actions in the Preconception Context

45 of 64 (70%) ARC were not pregnant when they received their results. Of these, 62% (28/45) responded that they pursued or planned to pursue alternative reproductive options, either IVF with PGD (n=22) or prenatal diagnosis (n=6). None selected the other options: gamete donation, adoption, no longer planning to have children. 29% (13/45) responded that they were not planning to pursue any alternative options based on the results. The remaining 9% (4/45) selected “other” and/or provided responses that were not indicative of a clear future direction. Of these 45 ARC, 31 had carrier screening as part of a fertility work-up. All ARC who did not pursue or plan to pursue alternative options carried moderate severity conditions (achromatopsia OMIM #262300, alpha-1 antitrypsin deficiency OMIM #613490, and *GJB2*-related DFNB1 nonsyndromic hearing loss and deafness OMIM #220290) or biotinidase deficiency (OMIM #253260), classified as a severe condition (see Discussion about genotype-phenotype spectrum in this condition).

### Qualitative Analysis of Decision Making

Respondents had the opportunity to provide free-text responses. Those who were not pregnant at the time of screening were asked about their reproductive choices/plans and presented with the opportunity to respond to the prompt “What factors influenced your decision?” ARC who were pregnant at the time of screening and who did not elect to pursue prenatal diagnosis were asked “What were some reasons you chose not to pursue prenatal diagnostic testing?” Those same respondents were asked about their reproductive choices/plans in future pregnancies now that they had received ECS results and then invited to respond to “What factors influenced your decision?” Thematic analysis was performed on these short, free-text responses. Some responses contained more than one theme.

Seventeen free text responses discussed reasons an ARC would choose not to pursue alternative reproductive options, with the dominant theme (14/17) being disease severity. These ARC indicated that they did not perceive the condition to be serious enough to warrant a change in reproductive planning:

> *“The carrier screening results we received were not linked with (in our perception) substantial pain or suffering a child might experience, and therefore not worth trying to prevent through an alternative conception/adoption option.”* (Preconception; *GJB2*-related DFNB1 Nonsyndromic Hearing Loss and Deafness)

Others perceived a low risk of an affected child with some referencing the risk from a low penetrance or mild allele. One referenced the risks of miscarriage inherent to prenatal diagnosis and another the cost of alternatives like IVF.

Of 21 ARC who provided free-text responses with their reasons for choosing to pursue alternative reproductive options, the dominant theme (11/21) also regarded severity. In this analysis desire for a healthy child or a child without the disease were coded as a subtheme of disease severity.

> *“Symptoms and severity of the condition, if inherited and the desire to not have an affected child.”* (Preconception; Hb Beta Chain-Related Hemoglobinopathy)

Others referenced the risk or chance of an affected child, indicating that they perceived the risk to be high and/or were unwilling to take the risk. Some respondents explored the choice to pursue IVF with PGD as a way to avoid the potential for the emotional pain of terminating a wanted pregnancy after prenatal diagnosis. Several indicated that they were already considering IVF and chose to add PGD after receiving the results to improve chances for a healthy pregnancy.

## DISCUSSION

The data demonstrate that carrier screening results affected clinical decision making for the majority of ARC. Couples who were carriers of a disease classified as profound or severe were significantly more likely to take action based on the results than those who were carriers for a moderate condition. The qualitative analysis of participants’ open text responses describing their decision-making rationale corroborated the quantitative analysis: the majority of responses concerned perceived disease severity. Inclusion in professional society guidelines also did not affect the likelihood of changed reproductive decisions.

This study also offers insight into the effect of incomplete penetrance and variable expressivity on reproductive decision making. Only 2 of 9 ARC in our data carrying biotinidase deficiency (BTD) reported changing their reproductive actions (2 provided unclear responses). The sharp contrast to other severe conditions raised the question of why the difference existed. While complete BTD is classified as a severe condition, partial deficiency (10-30% enzyme activity) is associated with a mild presentation. One variant, p.Asp444His (D444H, HGVS NC_000003.11:g.15686693G>C) occurs at high frequency (allele frequency of 3.94% = carrier frequency of 7.6% among non-Finnish Europeans in ExAC)^16,17^. It is a common cause of partial BTD when combined with a classic mutation, and is associated with approximately 50% enzyme activity in homozygotes 18, similar to the asymptomatic carrier state. While participants were asked to report the condition that they were carrier couples of, they were not asked to report the specific variants, making it impossible to directly ascertain which ARC carried D444H.

However, a review of the records of the original ARC invited to the study (including both respondents and nonrespondents) revealed that in 53 / 57 BTD ARC (93%) both members of the couple were D444H carriers. Our result that 7 of 9 BTD ARC did not report action taken based on this result is consistent with the hypothesis that all such ARC were D444H double-carriers (p=0.19). Thus, our results suggest that ARC appropriately considered their particular BTD results as a moderate condition. Notably, this indicates that detailed genetic counseling or laboratory-physician communication was effective, as the nuances of the particular test result seem to have been handled differently from the overall disease severity. This finding illuminates the variability of genetic conditions and may demonstrate a need for additional granularity in disease classification based on known genotype/phenotype correlations.

Questions have been raised regarding the ability of providers and patients to understand a wide variety of rare diseases sufficiently to make decisions based on their carrier status for one of them.^4^ One common caution regarding ECS is the overwhelming amount of information needed to counsel on each condition on the panel.^3^ Based on the clear differential in reproductive decisions between profound/severe and moderate conditions presented here, providers and patients understood the relative level of severity and impact of the various diseases identified by ECS. Moreover at least in the case of BTD, the specific genotype/phenotype correlations were also considered for their individual results.This may also be a result of the high uptake of post-test genetic counseling: 95% of ARC (61/64) pursued genetic counseling after receiving their results.

### Limitations of Study

In any survey research study where only a subset of eligible participants respond to the invitation, the data is not based on a random sample and may be affected by response bias. Accuracy may be limited by participants’ memory, interest in sharing their full experiences, and levels of medical literacy. Couples in the preconception setting were asked what options they pursued or planned to pursue after receiving their carrier screening results. Planned behaviors may not correlate to future actions.

Because of the limitations of survey response data and the limited offering of ECS outside of the fertility treatment context, caution may be needed in trying to apply the results of this study to the general population. Our sample was highly educated and high income, with many derived from infertility settings.

Future research should differentiate between the mutations carried by these couples and the probable genotype/phenotype correlations that could affect disease severity.

## CONCLUSION

This study reports the reproductive decisions made by ARC after receipt of ECS results to evaluate and demonstrate the clinical utility of ECS. Not only did the majority of ARC identified through ECS alter their reproductive decisions based on these results, but there was no significant difference between the rate of action between severe and profound diseases currently recommended by professional societies and those not yet included in screening guidelines.

## ACKNOWLEDGMENTS

The authors would like to thank Caroline Lieber, Janey Youngblom, and Kaylene Ready for their contributions to this research, as well as the couples who volunteered to participate in the survey.

